# An ancient plant symbiotic fungus with distinct features identified through advanced fluorescence and Raman imaging

**DOI:** 10.1101/2025.05.22.655410

**Authors:** Christine Strullu-Derrien, Raymond Wightman, Liam Patrick McDonnell, Gareth Evans, Frédéric Fercoq, Paul Kenrick, Andrea C. Ferrari, Sebastian Schornack

## Abstract

Mycorrhizal associations between fungi and plants are a fundamental aspect of terrestrial ecosystems. The occurrence of mycorrhizae is now well established in c. 85% of extant plants, yet the geological record of these associations is sparse. Fossil evidence from early Paleozoic terrestrial environments provides rare insights into these ancient symbioses, but imaging micro-scale fossil fungi in surrounding plant tissues is challenging. We employed brightfield microscopy, confocal laser scanning microscopy (CLSM), fluorescence lifetime imaging microscopy (FLIM), and Raman spectroscopy to investigate a newly identified endomycorrhizal fungus within a 407-million-year-old plant from the Windyfield Chert. An exquisite preservation and our advanced imaging approach enabled the description of the fungal structures to unprecedented detail. The fungus, *Rugosomyces lavoisieriae* gen. nov., & sp. nov., exhibits structural features resembling modern Glomeromycotina arbuscular mycorrhizal fungi. Combining CLSM with FLIM provides extra information by separating features based upon fluorescence lifetime. Raman spectroscopy indicates that fungal arbuscules and plant water-conducting cells have undergone geological alterations, resulting in a similar chemical composition. These findings enhance our understanding of ancient plant-fungal symbioses and demonstrate the potential of advanced imaging techniques in paleobotanical research.

**Significance statement:** Most living plants form mycorrhizal associations with fungi, but how this ancient evolutionary relationship developed is poorly understood. Exceptional fossil materials preserve direct evidence of endomycorrhizae, but imaging micro-scale fossil fungi inside plant cells is challenging. Here we use confocal microscopy combined with fluorescence lifetime imaging and Raman spectroscopy to study a new endomycorrhizal fungus inside a 400-million-year-old plant. We document structures resembling modern Glomeromycotina. We show that the residual organic matter of the fungus and the plant cell walls has a very short fluorescence lifetime, which provides a new means of imaging these fossils. Raman spectroscopy indicates that both arbuscules and water-conducting cells were geologically altered by their diagenetic history and that they now have a similar chemical composition.

## Introduction

Fungi and plants have a lengthy record, dating back to the early Devonian and they have interacted with each other in diverse ways for much of the history of life on land (1) (Berbee et al., 2020). The fossil record provides rare insights into the evolution of fungi as symbionts and pathogens of plants as well as crucial evidence for their roles as saprotrophs (e.g., 2–4). The oldest geological evidence for the endomycorrhizal symbiosis comes from 407-million-year-old cherts at Rhynie in Scotland (UK) (5, 6). The cherts formed as hydrothermal waters rich in silica spread across a wetland community entombing the organisms and preserving them in exquisite detail. The fossils within the cherts are studied typically in petrographic thin sections using brightfield microscopy (2, 6). Despite their frequently excellent preservation at Rhynie, fungi and other microorganisms are challenging to document due to their micrometer-scale size and to the limitations of brightfield microscopy of small organisms set within the chert material. We applied confocal laser scanning microscopy (CLSM) to the cyanobacterial (7), algal (8), protist (9), and fungal (10–12) components of this system to bring structures of a few micrometers into sharper focus through optical sectioning, and to reconstruct organisms and their interactions in three dimensions. Recently CSLM has also been used by MacMahon el al. (13) for describing another type of cyanobacteria.

The possibility that Rhynie plants hosted endomycorrhizae was first suggested by Kidston and Lang (14) in their original descriptions of the fossil fungi, and this idea was later developed by Boullard and Lemoigne (15). The first compelling evidence for arbuscules came from the sporophyte of *Aglaophyton majus* (16) in a fungus attributed to Glomeromycota (17) and later also noted in its young gametophyte (18). Further evidence of intracellular coils and arbuscules was reported in the plant *Horneophyton lignieri*. Two distinctve types of fungus were observed within different tissues of this plant (19) indicating simultaneous colonization by species attributable to both Glomeromycotina and Mucoromycotina.

Here, we combined CLSM with fluorescence lifetime imaging (FLIM) to investigate the fine structure of a new arbuscular mycorrhizal fungus observed within the aerial axis of the early land plant *Aglaophyton majus*. The material comes from the Windyfield chert, a new fossiliferous deposit discovered in 1989, about 700 m from the original Rhynie chert site (20). Unlike traditional fluorescence microscopy, which creates an image based on fluorescence emission intensity (21), FLIM is a technique that measures the time a fluorophore remains in an excited state before emitting a photon (22). Confocal-FLIM uses a pulsed laser and time-correlated single photon counting detection that gives an output of fluorescence decay, with a fast decay indicating short lifetime components, and slow decay indicating long lifetimes (22, 23). Lifetime changes are sensitive to the immediate changes in the environment of the fluorophore and its composition (23) and structure (24). This sensitivity was demonstrated in living plants, with changes in fluorescence lifetime of lignin, measured by confocal-FLIM, corresponding to small changes in the chemical structure of lignin, measured by Raman microscopy (25, 26). Because of the high sensitivity (measured with temporal resolution of picoseconds of FLIM to the chemical environment, here we use fluorescence lifetime as a means of imaging the fossilized remains of fungi and plants and of characterizing the nature of their interaction. Raman spectrometry is then employed to investigate the carbon framework of the fossils. Our results show that these are promising techniques for distinguishing between chemically different organic and mineral backgrounds.

## Results

We report the identification of fossil structures representing an endomycorrhizal fungus with structural features distinct from those of previously known fossil representatives. *Rugosomyces lavoisieriae* Strullu-Derrien & Schornack *sp. nov*. was found in fossilised axes of the plant *Aglaophyton majus* within the Windyfield Chert, dated to the Lower Devonian (27). The holotype specimen is preserved in slide no. NMS G.2022.11.48.1 at the National Museum of Scotland, Edinburgh (Fig. 1).

**Figure 1.**
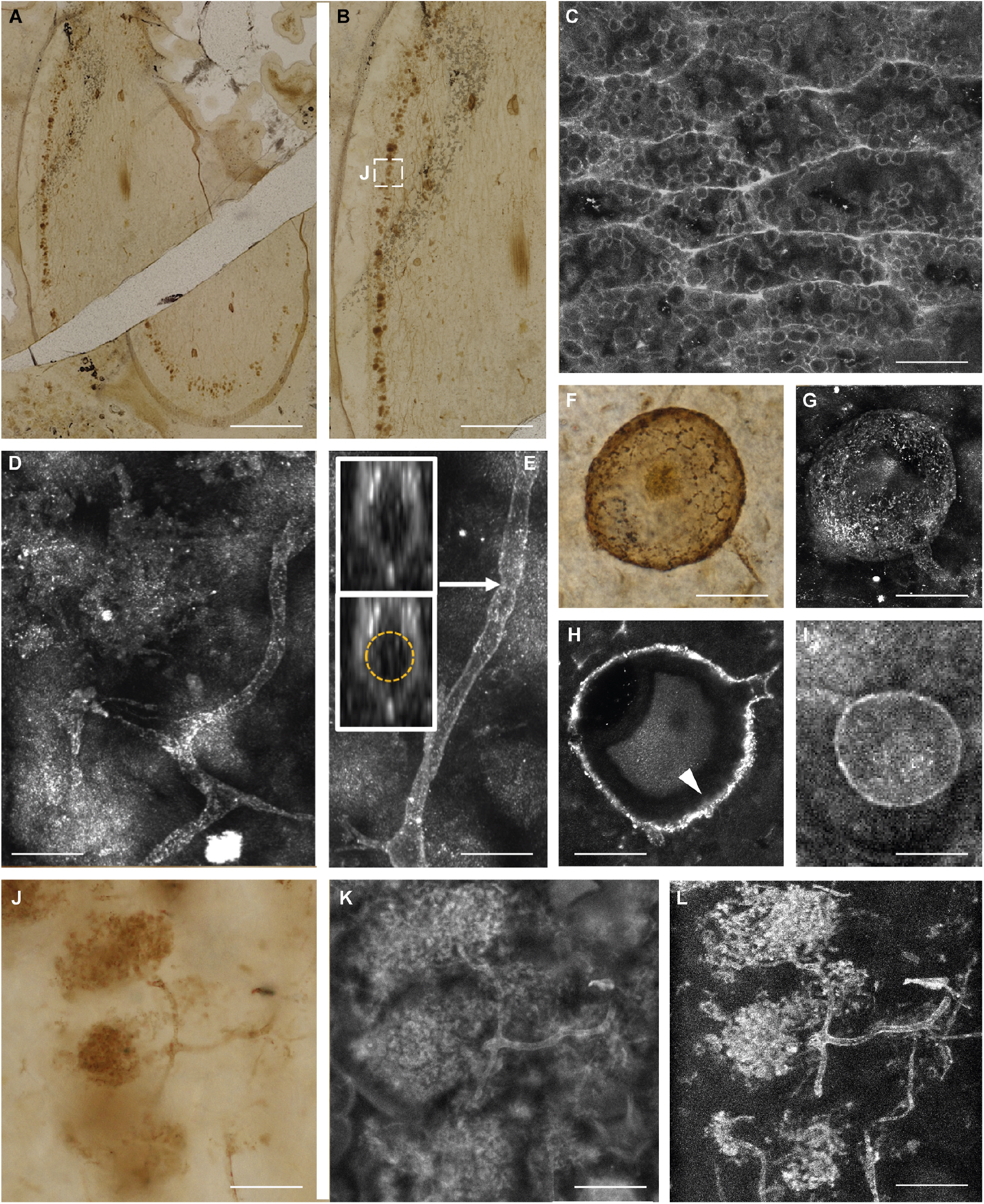
Arbuscular mycorrhizae in *Aglaophyton majus* and comparative microscopy. A, B Slightly oblique transverse section of the plant axis showing the fungal arbuscular zone at the perphery of the inner cortex (white box) and the central strand of water-conducting cells (brightfield microscopy). C. Water-conducting cells of the vascular strand showing the vesicular structure (CLSM spectral). D. H-branching in hyphal development (CLSM). E. Hypha with pseudoseptum. Sections through the hypha (insets) show that there is no internal septum. (CLSM). F. Spore showing rugose surface (brightfield microscopy). g. Maximum projection image of a fungal spore (CLSM). H. Single orthoslice image showing the structure of the wall and the septum on the branching hypha (CSLM & Airyscan). I. Vesicle and branching hypha (CLSM). J-L. Comparative microscopy: arbuscules in brigthfield microscopy (J), CLSM (K) and FLIM – short (0.3 ns) lifetime (L). Scale bars: 1,1 mm; (A); 0.7 mm (B); 25 µm (C, H); 24 µm (D); 18 µm (E); 45 µm (J); 30 µm (F, G, K); 38 µm (l). Slide n° NMS G.2022.11.48.1. See also the original datasets: https://doi.org/.10.5281/zenodo.15194427 (59).

### 1. Fluorescence lifetime imaging distinguishes fungal from plant structures

We image both fungus and tissues of the host plant using brightfield (Fig.1 A, B, F, J) and confocal laser scanning microscopy (CLSM) (Fig. 1C–E, G–I, K, L) and fluorescence lifetime imaging microscopy (FLIM) (Fig. 2). CLSM reveals key features of the hyphae (lack of septum, Fig. 1E), fungal spores (detail of the spore wall and subtending hypha, Fig. 1G, H), vesicles (Fig. 1I) and arbuscules (Fig.1J–L). CLSM-based imaging outperforms brightfield microscopy by improving image clarity (Fig. 1J versus Fig. 1K). For plant tissues, CLSM reveals the vesicular structure of the water-conducting cells of the host plant *A. majus*, a well-preserved feature of this tissue (Fig. 1C).

**Figure 2.**
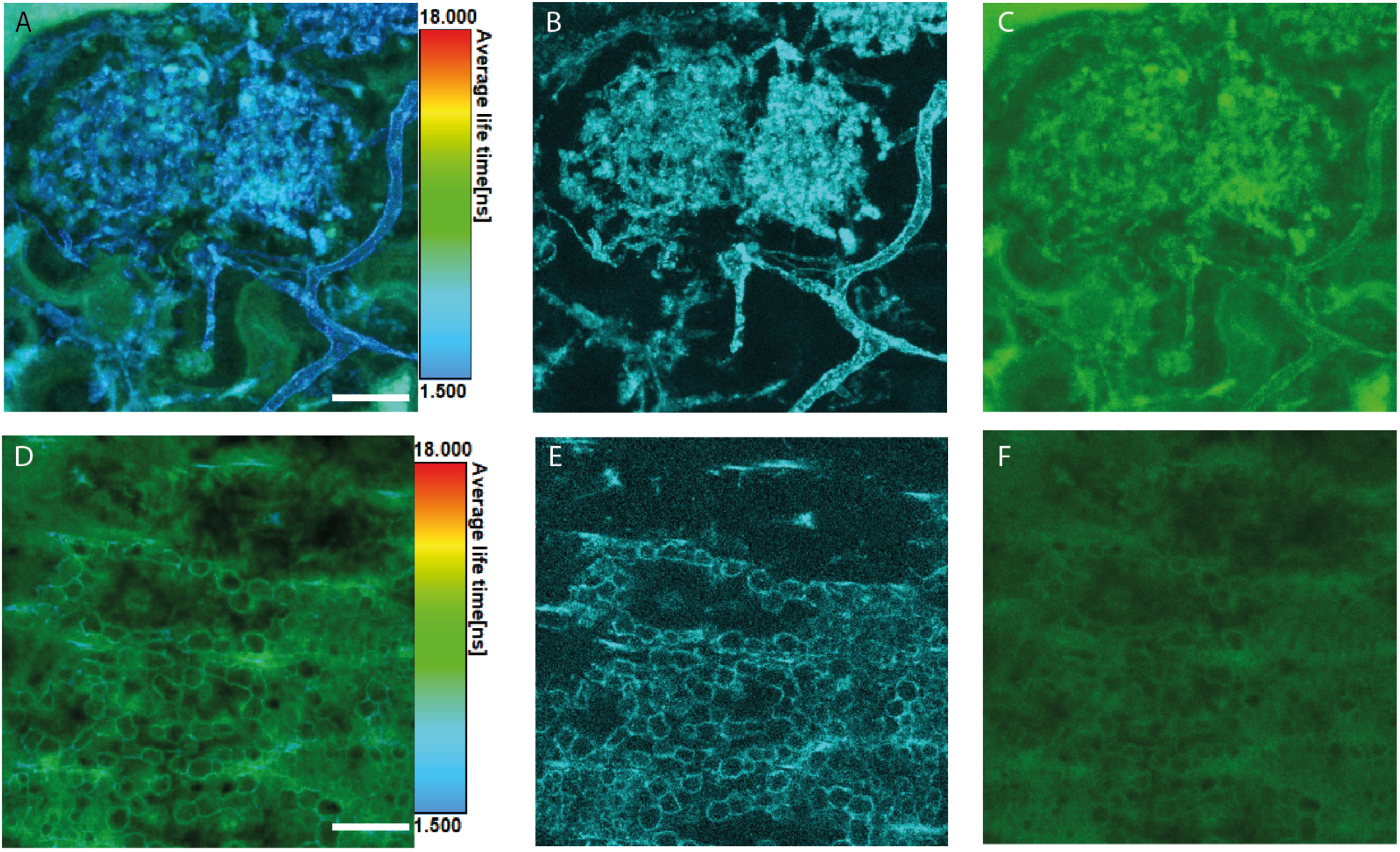
FLIM analyses. A. Maximum projection of a 23 slice Z-stack showing relatively short average lifetime of autofluorescence of the arbuscules (blue/cyan) compared to areas of the plant devoid of fungal structures (green). B. Maximum projection of the short lifetime that represents the arbuscules only. C. Maximum projection of the long lifetime representing plant-plus-arbuscules. D. Single optical section through the central strand of the aerial axis of *Aglaophyton majus* showing average lifetime of autofluorescence. Same optical section showing the short (picosecond) lifetime (E) and long (nanosecond) lifetime (F). The short lifetime gives high signal in the ovoid structures (vesicles) within the water-conducting cells. Scale bars: 20 µm. See also the original datasets: https://doi.org/.10.5281/zenodo.15194427 (59).

FLIM of a fungal arbuscule located in a background of plant material preserved in a silica matrix (i.e. chert) returns average fluorescence lifetimes that emphasize different features of the specimen (Fig. 2). A short average fluorescence lifetime of 0.3 ns separates the fungus (arbuscule and other hyphae) from background (Fig. 2B). Longer average lifetimes >6 ns give signals from both fungus and background (Fig. 2C). Combining both shorter and longer lifetimes highlights where the background contributes uniquely to the longer fluorescence lifetime (Fig. 2A). Similarly, FLIM of the plant water-conducting cells in a background silica matrix, but without obvious fungi, returns average fluorescence lifetimes that pick out different features of the specimen (Fig. 2 D–F). The short 0.3 ns lifetime gives a clear signal from the vesicles of the conducting tissue (Fig. 2E). In contrast to the arbuscule, the longer >6 ns lifetime provides a much more uniform signal of lower intensity from plant and background and there is little structure in the image (Fig. 2F).

Compared to other light microscopy methods, the imaging of fungus and host plant using CLSM gives better definition of cell structures due to the improved axial resolution that is inherent to CLSM. Combining CLSM with FLIM provides extra information by separating features based upon fluorescence lifetime (Supplementary Movie). By observing just the short 0.3 ns fluorescence lifetime, a clearer image that eliminates background can be obtained for both fungal arbuscules and vesicles of the plant water-conducting cells. This yields greatly improved signal-to-noise ratio, and the residual organic-derived carbon in the fossil is the source of this short lifetime fluorescence. The sources of the longer (>6 ns) lifetime fluorescence encompass both fungal hyphae and possibly also plant cell derived materials. The more uniform background signal in both the arbuscular zone and the pro-vascular cylinder indicates that the silica matrix might also be contributing to the longer lifetime.

### 2. Raman investigation of the fluorescent carbonaceous material of the fungal and plant cell walls

To probe the nature of the organic signal underpinning the FLIM results in figure 2, the arbuscules of the fungus and the water-conducting cells in the vascular system of the host plant are analysed using Raman spectroscopy (Fig. 3A-H).

**Figure 3.**
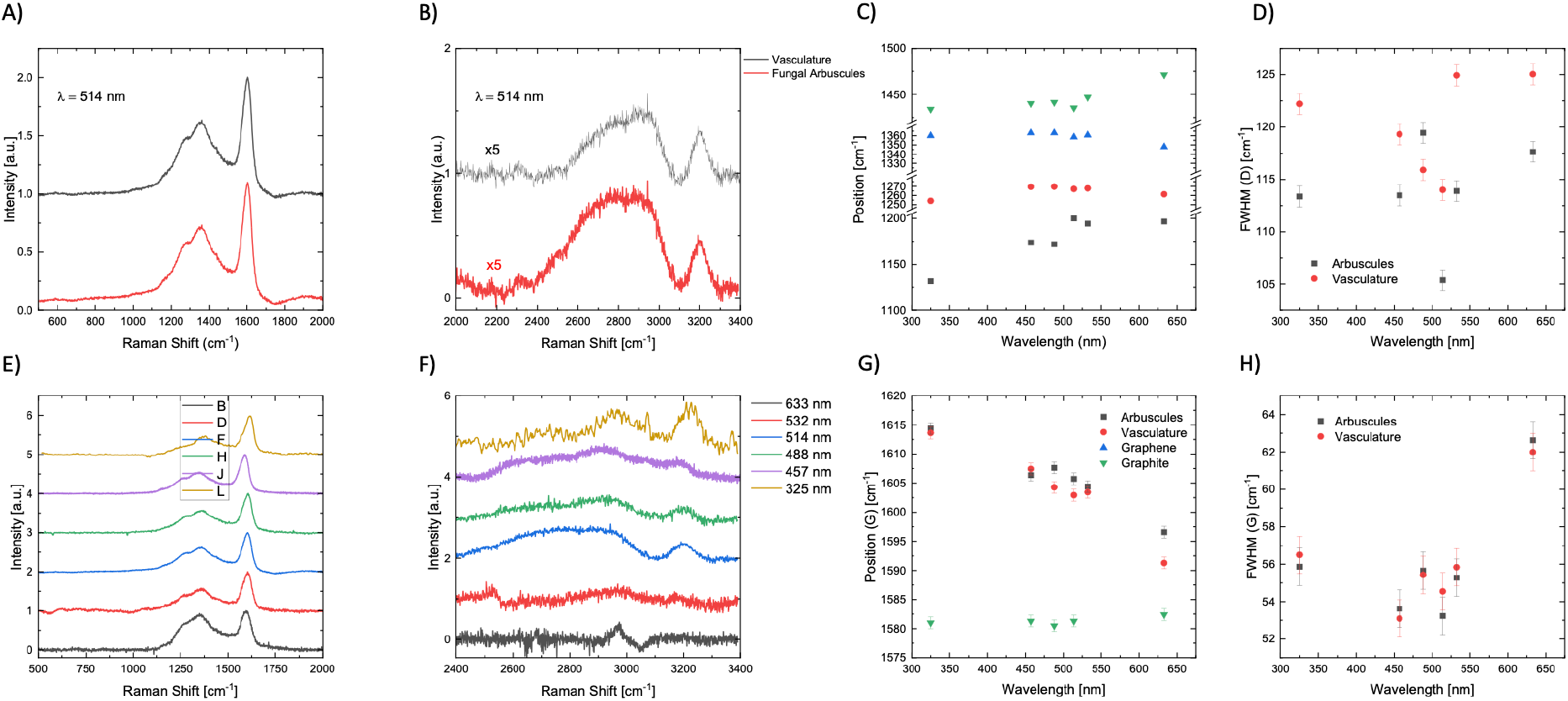
Raman analyses. A, B. Raman spectra for vascular strand (black) and arbuscules (red) measured at 514nm. E, F. Raman spectra acquired at multiple wavelengths for the arbuscules. C, G. Pos(D), Pos(G) and of subbands∼ 1266 and 1450 cm^-1^ at 514nm, as a function of wavelength. D, H. FWHM (G) for both fungal arbuscules and vascular strand.

Raman spectra were taken for both the fungal arbuscules and the water conducting cells as shown in figures 3A and 3B. For both samples the spectra are characterized by ∼ 4 peaks between 1000 to 1650 cm^-1^. The spectra acquired at 325, 457 and 488 nm also show several peaks between 2800 and 3200 cm^-1^ in both samples (Fig. 3E, F). Peaks ∼1358 and 1608 cm^-1^ are assigned to the D and G peak respectively and arise due to the presence of sp^2^ carbon (28, 29). The UV spectra in figure 3F have peaks ∼2960 and 3220 cm^-1^, assigned to C-H stretching and to the 2D’ respectively (30). Analysis of the D and G peaks including their positions, Full Width at Half Maximum (FWHM) (Fig. 3D, H), intensity ratios, dispersion derived from the peak positions in figures 3C and 3 G and henceforth referred to as Disp (i.e. the shift in frequency as a function of excitation energy) can be used to determine the nature of the underlying sp^2^ carbon. FWHM(G) for both fungal arbuscles and water conducting tissue is ∼56 cm^-1^ at 514 nm, consistent with nanocrystalline sp^2^ domains (28, 31). The peaks ∼ 1266 and 1433 cm^-1^ (at 514 nm) are consistent with those found in carbon chains (28, 32, 33), possibly derived from the transformation of precursor compounds in the arbuscules and water conducting cells such as lignin, chitin or cellulose.

For the fungal arbuscules as well as the water conducting cells Disp(D) is ∼4.1 (8.2) cm^-1^/eV and Disp(G) is ∼8.3 (10.3) cm^-1^/eV, consistent with both regions containing similar sp^2^ carbon (28, 33). The much lower Disp(D) with respect to graphite (28, 33), means that sp^2^ domains only span a limited distribution in sizes, and do not combine to form a long-range graphitic lattice.

The intensity ratio of the D and G peaks, I(D)/I(G), can be used to estimate the mean size L_a_ of the sp^2^ clusters (34, 35). For the fungal arbuscules and water conducting cells we have I(D)/I(G) ∼0.47-0.6 at λ=514nm, implying a mean length of the sp^2^ clusters∼1.1-1.2 nm (35-37). These data suggest that the investigated fossilized fungal and plant material has undergone diagenetic alteration, resulting in the formation of nanocrystalline carbon structures.

### 3 A morphologically distinct endomycorrhizal fungus colonizing the plant *Aglaophyton majus*

#### 3.1. Systematics

Kingdom – *Fungi* R.T. Moore.

Phylum – *Mucoromycota* Doweld, emend. Spatafora & Stajich.

Subphylum – *Glomeromycotina* (C. Walker & A. Schüßler) Spatafora & Stajich, subphylum and *stat. nov*.

Class – *Glomeromycetes* Caval.-Sm.

Order – *Incertae sedis -* Strullu-Derrien & Schornack.

Genus – *Rugosomyces* Strullu-Derrien & Schornack *gen. nov.*

Etymology – *Rugoso* refers to the nature of the spore wall; *myces* is the latin word for fungus. Genus Diagnosis – Fungus with aseptate intercellular hyphae with H branching, spores, vesicles and intracellular arbuscules. Differs from other fungi in the Rhynie chert in the following ways: spore wall possibly bilayered, with a relatively thick, rugose wall; smaller diameter of the branching hyphae; hypha terminating in a spore are either smaller or larger than other hyphae; the ratio of spore diameter to hyphal width is smaller; the spores are comparatively either smaller or larger.

Species – *Rugosomyces lavoisieriae* Strullu-Derrien and Schornack *sp. nov.*

Etymology – In honour of Marie-Anne Paulze de Lavoisier (1758-1836) who was a collaborator of her husband Antoine Laurent de Lavoisier and was his laboratory assistant. She translated Richard Kirwan’s ‘Essay on Phlogiston’ from English to French which allowed her husband and others to dispute Kirwan’s idea. The scientific collaboration of this husband-wife team is perhaps unique among the giants of respiratory physiology (38).

Species diagnosis – Branched hyphae 3.5–6.5 µm in diameter; unbranched hyphae 7 µm in diameter when terminating in a spore and 4 µm in diameter when terminating in a vesicle; basal stalk of the arbuscules 3.5 µm in diameter. Spores globose, up to 74 µm in diameter, with a 3.1 µm thick rugose and possibly bilayered wall. Subtending hypha closed by a septum. Vesicles from globose (up to 39 µm in diameter) to elongate (41 µm wide, 50 µm long).

Holotype – specimens in slide no. NMS G.2022.11.48.1 at the National Museum of Scotland, Edinburgh. Fig. 1.

Locality – Rhynie, North-West of Aberdeen (Scotland): Windyfield Cherts Unit (39). Age – Lower Devonian (407.1 ± 2.2 Ma) (27)

Mycobank (40, 41) nos: (will be provided after acceptance of the manuscript).

#### 3.2. Description of fungus and its known distribution in the plant

*R. lavoisieriae* is observed in an oblique transverse section of an axis of the fossil plant *Aglaophyton majus* (Fig 1A, B). The outlines of epidermal cells are preserved in the cuticle. The distinctive cylinder of cells shows a vesicular structure in the center of the axis (Fig. 1C; Fig. 1A in Supplementary Appendix). This corresponds to the inner zone of thin-walled water-conducting cells that is typical of the vascular system of *A. majus*, the vesicles being a diagnostic feature of this tissue (42, 43).

Fungal colonization occurs in a circumferential continuous zone (Fig. 1A). Fungal structures such as hyphae, spores, vesicles, and intracellular arbuscules differ in their relative abundance in the different zones of the plant axis. The inner cortex is dominated by intercellular hyphae and surrounded by a zone in which arbuscules develop within turgescent cells (see Fig.1B in Supplementary Appendix), and which also contains hyphae, vesicles and spores; the outer cortex is devoid of fungal colonization (Fig. 1B). Hyphae form H branching (Fig. 1D). Detailed investigation of the hyphae in 3D reveals pseudo-septa that seem to be bends in the hyphal wall rather than true septa (Fig. 1E), thus all hyphae are aseptate except for the point of attachment of the spore in which the hyphal lumen is closed by a septum (Fig. 1H). Hyphae are variable in width: branched forms range from 3.5 µm to 6.5 µm in diameter; unbranched forms are 7 µm in diameter when terminating in a spore and 4 µm in diameter when terminating in a vesicle (Fig. 1F-I). Spores are globose (up to 74 µm in diameter, possibly bilayered although layers are not easily discernible (Fig. 1H), with a thickened (3.1 µm thick) rugose wall (Fig. 1F–H). Vesicles, corresponding to hyphal swellings, are of varying shape from globose (up to 39 µm in diameter) to elongate (41 µm wide, 50 µm long) (Fig. 1I). Intracellular arbuscules develop at the top of branched hyphae 3.5 µm in diameter (Fig. 1J-L). Hyphae of this type have not been observed in proximity and external to the plant axis, so we were unable to illustrate the point of penetration.

#### 3.3. Comparison with endomycorrhizal Glomeromycotina at Rhynie

*Rugosomyces lavoisieriae* differs from other endomycorrhizal Glomeromycotina at Rhynie primarily in its spores, vesicles, and its ratio of spore to subtending hyphal diameter. *Rugosomyces lavoisieriae* can be distinguished from *Glomites rhyniensis* (17) primarily on the size and structure of the spores and vesicles and on the dimensions attained by hyphae in the inner cortex. The hyphae of both species are aseptate, but occasional septation of narrow thin-walled hyphae was noted in *G. rhyniensis*. It is difficult to be sure that these are true septae because they might be a taphonomical artefact like the pseudoseptae that we observed in *R. lavoisieriae* (Fig. 1E). H-branching is a feature of the hyphae in both species, but the hyphae in the inner cortex of *R. lavoisieriae* are much narrower (3.5–6.5 µm) than those of *G. rhyniensis* (8–14 µm). The spores of *G. rhyniensis* have a smooth, thick (6 µm), multilayered-wall with an inner layer that is continuous with the subtending hypha, whereas the spores of *R. lavoisieriae* have a thinner (3.1 µm), rugose, possibly bilayered-wall with a septum on the subtending hypha. The spores of *G. rhyniensis* are globose to elongate and 50–80 µm in diameter, whereas those of *R. lavoisieriae* are more globose but of similar maximum size (∼74 µm). However, the spore category of *G. rhyniensis* also encompasses vesicles, which in *R. lavoisieriae* are elongate or globose but much smaller structures (41 x 50 µm; up to 39 µmin diameter). The arbuscules of *G. rhyniensis* and *R. lavoisieriae* are indistinguishable: both developed from narrow hyphae (∼3.5 µm wide) giving rise to 2–3 major branches that led to clusters of much branched narrower hyphae.

*R. lavoisieriae* can be distinguished from *Palaeoglomus boullardii* (19) primarily on the structure of the spores. *R. lavoisieriae* has spores with a thicker (3.1 µm), rugose wall, and a septum on the subtending hypha, whereas those of *P. boullardii* are thinner (1.5 µm), smooth-walled and have a subtending hypha that is continuous with the spore wall and closed by the invaginated inner layer of the spore wall. The ratio of spore to subtending hyphal diameter is higher in *P. boullardii* (15-18:1), and *G. rhyniensis* (10-18:1) than in *R. lavoisieriae* (10:1).

We have not found report of a similar fungus from deposits of this age and therefore describe it as new here.

#### 3.4. Comparison with modern mycorrhizae

The intracellular fungal colonization in the aerial axis of *A. majus* by *R. lavoisieriae* resembles the colonization by *Glomeromycotina* of extant thalloid liverworts, hornworts, lycophytes and ferns (19, 44-48). Similarities include form, size and structure of the spore wall, H connections between parallel strands, intra-axial vesicles and arbuscules. Furthermore, spores have a single hyphal attachment that persists, which is also a feature of extant *Glomus*. Because Devonian fossil plants are evolutionary and structurally closer to extant bryophytes and lycophytes, we argued that comparisons with the latter rather than with gymnosperms and angiosperms are much more appropriate when interpreting the anatomy of early land plant–fungus symbioses (19).

## Discussion

Three species of endomycorrhizal fungi showing clear evidence of arbuscules have now been documented in two species of plant from Rhynie (17, 19, this paper). Our new species is the second species to be recorded in *Aglaophyton majus*. The first was *Glomites rhyniensis* (17). These two records come from two different stratigraphic units within the chert beds at Rhynie. *G. rhyniensis* was described from the Rhynie Cherts Unit, which is the main and best-known source of chert. *R. lavoisieriae* comes from the Windyfield Cherts Unit, which is a separate, smaller and highly localised near surface exposure located 700 m to the north-east of the main deposit (49). The Windyfield Cherts Unit is slightly younger stratigraphically than the Rhynie Cherts Unit (39), but they both belong to the same spore biozone and are therefore indistinguishable in terms of their geological age (50). The biota of the Rhynie and the Windyfield Chert Units are similar in terms of plant, protist and bacterial components, but there is a slightly greater richness of arthropods at Windyfield (20, 51). In terms of palaeoenvironment, the two chert units are also similar, but deposition at Windyfield took place closer to the centre of hydrothermal activity. Fayers and Trewin (51) noted in the caption of their figure 6G: “endotrophic mycorrhizae in the cortex of a *Ventarura* rhizome”, but no description was provided, and the image of fungal spores is not informative enough to support the conclusion that mycorrhizae were present. *R. lavoisieriae* is therefore the first documented endomycorrhizal fungus from the Windyfield Chert Unit.

The patterns of tissue colonization in *Aglaophyton majus* by *Glomites rhyniensis* (17) and by *Rugosomyces lavoisieriae* are broadly similar. Both developed an extensive network of intercellular hyphae in the inner cortex and a very distinctive zone of intracellular arbuscules in a narrow ring at the perimeter of the inner cortex. The colonization of the hypodermis (i.e., outer cortex) by *G. rhyniensis* differed from that of the inner cortex. In the hypodermis the hyphae were thick-walled and highly branched. Taylor *et al.* (17) also documented large (25 µm diameter) ‘extraradical’ hyphae forming cord-like structures. Neither hypodermal colonization nor extra-axial hyphae were observed in *R. lavoisieriae*. Comparison between *Rugosomyces lavoisieriae* and *Glomites rhyniensis* (above) shows that *Aglaophyton majus* formed symbioses with at least two distinct species of endomycorrhizal fungi that had similar general patterns of tissue colonization, but which differed primarily in the size and structure of their spores and vesicles and in the dimensions of certain hyphae.

Living plant cells and fungal hyphae have distinctly different chemical compositions. Fungal hyphae are rich in chitin whereas the plant water-conducting cells imaged were predominantly composed of cellulose but probably also contained a lignin component. The original chemistry of the cell walls becomes altered during fossilization and diagenesis, but evidence from several sources indicates that taxon-specific and even tissue specific signals persist in the organic remains of the organisms at Rhynie. High resolution analysis of tracheid cell walls of the fossil plant *Asteroxylon mackiei* using carbon X-ray absorption near-edge spectroscopy (C-XANES) provided evidence of hydroxylated aromatic and aliphatic carbon in the inner and outer wall regions, which was interpreted as reflecting the distribution of lignin and structural polysaccharides (52). The same method later showed that the conducting cells of *A. majus* and *Rhynia gwynne-vaughanii* contained less aromatic carbon than those of *A. mackiei* (53). The same oxygen-bonded carbon (i.e., C–O, C═O and O–C═O) was detected in *R. gwynne-vaughanii* using time-of-flight secondary ion mass spectrometry (ToF-SIMS), but the aliphatic/aromatic ratio was not seen to vary greatly across tissue systems (54).

The Raman spectra of our sample provide information about the organization of the aromatic skeleton, mostly about sp^2^ bonds (28, 33, 35). The spectra show that the carbonaceous materials of the arbuscules and the water-conducting cells are geologically altered by their diagenetic history and that they now have similar composition (Fig. 3a-h). The G and D peaks and sub bands at ∼1266 and 1450 cm^-1^, along with their dispersive behaviours indicate the presence of sp^2^ clusters ∼1.1nm in size as well as carbon chains and C-H bonds. The mean length of the sp^2^ clusters is in broad agreement with High-Resolution Transmission Electron Microscopy (HRTEM) results from Delarue et al. (55) for another Rhynie Chert fossil where the length scale of carbon clusters was found to be ∼0.55 nm. Loron et al. (56) performed ATR-FTIR studies on Rhynie chert microorganisms. They found absorption bands of organic matter in the interval 3000–2800 cm^−1^ and 1800–1400 cm^−1^ that they interpreted as characteristic absorptions for different CH, C=O, COOH, and nitrogen-moieties. They suggested that these organic groups represent the fossilization products of biomass that was originally dominated by lipids, proteins, and sugars. Then they transformed the intensity of informative organic bands (as revealed by the multivariate analyses) for each specimen (excluding the plant spores) into a matrix and conducted a discriminant analysis. The authors considered that the organic matter comprising Rhynie chert fossils is original and that the differences they observed are variations in the original precursors of the fossil organic matter, modified through diagenesis. However, one of the objects they identified as fungi appears to be a cluster of plant rhizoids (see Fig. 1c in 56, supplementary information). Another study was performed on Rhynie chert carbonaceous material by Raman spectrometry which indicated that the carbonaceous material has experienced advanced diagenesis (57), however regarding the differences noted between fungi and plants and within plant tissue systems, the results should be treated with caution because some of the identifications appear to be erroneous (see Supplementary Appendix).

In our specimens the residual carbonaceous materials of the fungus and the vesicles in the plant cell walls return a similar very short (0.3 ns) lifetime fluorescence. These structures are also the most resilient ones within the plant axis, and they have a similar brown colour under brightfield illumination. The explanation for the similar fluorescence lifetimes is that the chitin in the fungal arbuscules and the lignocellulose vesicles in the water-conducting cells have decomposed to simpler but similar chemical breakdown products as shown by the Raman analyses. This result can be compared to those obtained on the earliest known wood (from coeval deposits than the Rhynie Chert), which was anatomically exceptionally preserved. Transmission X-ray Microscopy-based X-ray Absorption Near Edge Structure and Transmission electron microscopy data performed on specimens preserved in 2D and 3D also show that the fossil wood was geologically altered by its diagenetic history and that the remaining organic matter, in both cases, has a similar chemical composition that does not reflect the original composition of the wood, even if a lignin source for the original compounds cannot be ruled out completely (58)

In conclusion, a combination of brightfield microscopy, advanced confocal-FLIM imaging and Raman analyses enabled the identification of a 407-million-year-old arbuscular mycorrhizal symbiosis in the Windyfield chert. An exquisite preservation and use of advanced imaging techniques enabled the description of carbonized fungal structures to unprecedented detail justifying a (re)-analysis of other fossil specimens using Confocal-FLIM analyses. The Raman analyses show that anatomical and chemical preservations may not be correlated, confirming previous results. It will be interesting to explore how abundant *R. lavoisierae* is within the Rhynie and Windyfield cherts, the extent to which it was able to colonize different tissues (e.g., rhizoids) or host plants, and to further explore the diversity of symbiotic associations in the Windyfield chert, thereby shedding new light on the diversity of plant-fungal interactions in the fossil record.

## Author contributions

C.S.-D. and S.S. conceptualized the study. C.S-D., S.S. and G.E. collected the data in brightfield microscopy. C.S.-D., F.F., S.S., R.W. and G.E. collected the confocal data; R.W. collected the FLIM data. L.P.McD. and A.C.F. performed the Raman analyses. All the authors analyzed their data. C.S-D. and P.K. wrote the original draft with the input of R.W., S.S., L.P.McD. and A.C.F. All authors contributed to the final version of the manuscript.

The authors declare no competing interest.

## Acknowledgements

The authors thank Andrew Ross from the National Museum of Scotland, Edinburg for rock supply and Callum Hatch for the preparation of the slide. We acknowledge funding by the Fondation ARS Cuttoli-Paul Appell/ Fondation de France to C.S-D (grant no. 00103178), by the Gatsby Foundation to S.S. (GAT3731/GLD), R.W. and G.E. (GAT3842), by the European Research Council and the UK Engineering and Physical Sciences Research Council (ERC Grants Hetero2D, GSYNCOR, EU Grant CHARM, EU Graphene Flagship, EPSRC Grants EP/Y035275/1, EP/V000055/1). The MNHN light microscopy facility (CeMIM, Centre de Microscopie et d’Imagerie du Muséum, MNHN Paris) is also thanked for providing access to the Zeiss 880 confocal scanning laser microscope.

This is a contribution to the Natural History Museum’s Evolution of Life Research Theme.

## Materials & Methods

The fossiliferous chert deposits at Rhynie are located in an outlier of Lower Old Red Sandstone situated about 50 km north-west of Aberdeen, Scotland, UK (60). Here, there are two distinct chert units that developed from their own centres of surface hydrothermal activity with hot springs and geysers. The main and best-known source of chert (Rhynie Cherts Unit) lies just below the surface of agricultural land (60). In 1989, a separate, smaller and highly localised near surface exposure of chert was found 700 m to the north-east of the main deposit (Windyfield Cherts Unit) (49, 51). The materials described here come from the Windyfield Cherts Unit, from which the quantity of material available for study is small and there are far fewer prepared thin sections.

Sinter deposition of both chert units originally took place in an intermontane basin on the southern margin of the palaeocontinent of Laurussia, with sediment accumulating on a low-energy alluvial plain (39). Palaeoenvironments range from terrestrial and vegetated sinter sheets to low-temperature pools and marginal aquatic settings. Silicification ensued from geothermal outwash of alkali-chloride hot springs at some distance from vents and probably at low temperature (<40-50°C) (51, 61). Plants were silicified close to their sites of growth and occasionally in growth position, together with their substrates.

A series of thin sections were prepared by Callum Hatch at the Natural History Museum London using the petrographic standard method. They were cut from a Windyfield chert block owned by the National Museum of Scotland, Edinburgh. Sections are c. 70 μm in thickness; they are mounted without glass coverslips, but the rock surface is highly polished. The results presented here are from slide slide no. NMS G.2022.11.48.1. The specimens were studied with a Nikon Eclipse LV100ND compound microscope at the Natural History Museum; depth of field was enhanced through z-stack montage. Digital light microscopy was also carried out on a Keyence VHX7000 at the Sainsbury Laboratory, Cambridge with overview images made up from merged tile scans.

For the analyses in CLSM, preliminary overviews and imaging for Fig. 1C and 1H were obtained at the MNHN light microscopy facility (CeMIM, Paris) using a Zeiss LSM880 confocal microscope and a plan Apo objective lense 40X1.30 NA – oil immersion objective lens. Samples were excited with the 561 nm laser line and auto-fluorescence signal was collected with either the Airyscan head using a 32 GaAsP detector array in super resolution mode or with the 32 channel GaAsP spectral detector. Images were recorded with pixel dimensions of 50-70 nm and 16-bit depth mode. For 1C, a 13 µm z-stack was acquired with steps of 0.5 mm. Images were processed in Zen Black software (Carl Zeiss) to deconvolve Airyscan images (Airyscan processing) and spectral images (Linear unmixing). For linear unmixing, the main autofluorescence spectrum was extracted using the “Auto Find” option in Zen Black.

Then CLSM imaging was performed at the Sainsbury Laboratory, Cambridge using a Leica SP8 X upright confocal microscope with a 561 nm laser and a HyD detector with an emission window of 633-791 nm. A 63x 1.4 NA oil immersion objective lens was used. Z stacks were acquired using the software optimised z-spacing.

Acquisitions on Fluorescence Lifetime Imaging microscopy (FLIM) were carried out on a Leica SP8-SMD upright confocal microscope equipped with Picoquant FLIM hardware running both LAS X and SymphoTime software. A 440 nm pulsed laser was used as excitation source at 20 MHz. The sample was viewed using a 63x 1.4 NA oil immersion objective. Detection of the signal used a SMD-HyD detector set to an emission window between 601 and 786 nm, photon counting mode. The laser power was altered to achieve a max count rate of 2000 cps. FLIM settings on the LAS X were 848 x 848 pixel format, unidirectional scanning, zoom 1.75 and with 70 FLIM iterations. Z stacks were acquired using LAS X software-optimised z-spacing. Average lifetime maps were generated using the FastFLIM calculations and displayed on SymphoTime. Curve fitting and extraction of images representing the discrete lifetimes was carried out on an offline SymphoTime workstation. The data follows a triple exponential decay with a very short picosecond range lifetime representing fungal structures only and a long nanosecond lifetime representing fungus plus plant. An intermediate low nanosecond lifetime represents low intensity background.

Raman spectra are acquired using Renishaw Invia and Horiba HR800 spectrometers using excitation wavelengths of 325, 457, 488, 514, 532 and 633 nm. For all wavelengths, the incident power is limited to <1mW to avoid both local heating and damage to the sample. Most spectra are measured using a x100 objective with a Numerical Aperture (NA)=0.95 resulting in a laser spot size∼1µm. The 325nm measurements use a x40 objective suitable for Near Ultra Violet (NUV) excitation. At each wavelength, the spectra are calibrated using the Si Raman peak at 520.7cm^-1^ (52). Spectra are also acquired away from the ROIs to remove background scatter and luminescence during baseline correction.

## Data availability

All confocal data collected and used in this study are deposited in the Zenodo repository under a Creative Commons Attribution 4.0 international license. https://doi.org/.10.5281/zenodo.15194427 (59). Slide no. NMS G.2022.11.48.1 is housed at the National Museum of Scotland, Edinburgh. Yves Candela can be contacted for access.

## Supplementary Appendix

### 1.1 Identification of *Aglaophyton majus* - the plant hosting the arbuscules

The identity of the large plant containing the arbuscular fungus was determined as *Aglaophyton majus* based on observations of axes in two consecutive thin sections made from the same block of chert (slides no. NMS G.2022.11.48.1 and no. NMS G.2022.11.48.2 at the National Museum of Scotland, Edinburgh).

The fungal arbuscules imaged with the confocal microscope are inside an axis in slide RW 14.1. The outlines of epidermal cells and two stomata are preserved in the cuticle. Few other plant cells are clearly distinguishable. At low magnification, there is a distinctive zone of brown bodies; this is the fungal arbuscular zone. These brown-coloured bodies of low opacity measure∼60–80 µm in diameter. The arbuscules are well identified by confocal and FLIM microscopy. They colonized turgescent cells (Fig. S1A) meaning that the plant was alive at the time of their formation.

The most distinctive tissue system that remains is a part of the vascular system in the centre of the axis. This is a cylinder of cells which corresponds to the inner zone of thin-walled water-conducting cells in the vascular system of *A. majus*. Further confirmation of this identity comes from high-resolution imaging with the confocal microscope (Fig. S1B). These cells contain numerous vesicles of varying size that are typically associated with a thin underlying cell wall. The vesicles are organised into chains that can partly occlude the cell lumen. The perimeter of the vesicles is somewhat angular. These distinctive features are diagnostic of the water-conducting cells of *A. majus* (see Remy & Hass, 1996; Plate IV fig. 5). Their origin and function are poorly understood. They have been variously interpreted, and more recently as part of the original cell wall structure (Remy & Hass, 1996; Edwards, 2003; Kerp, 2018). It is the first time that these vesicles have been observed in 3D.

**Figure S1.**
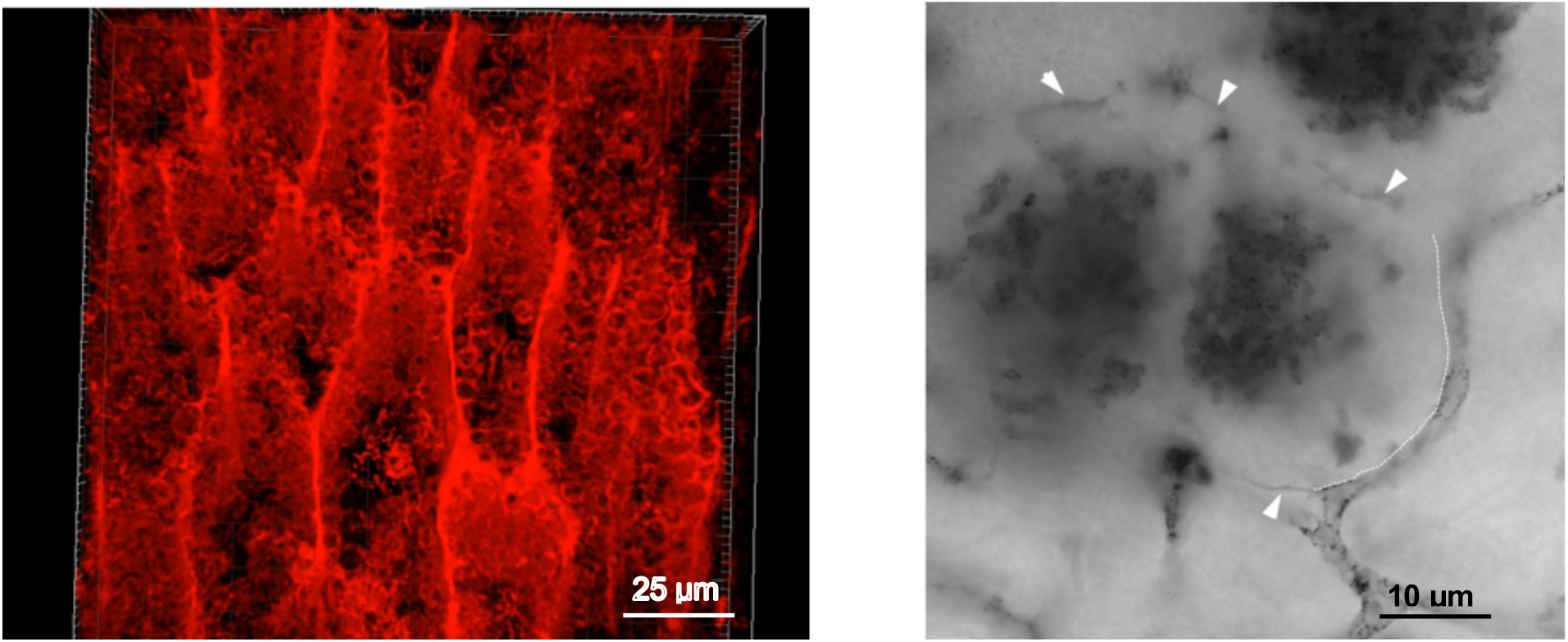
A. Vesicles in the vascular system. 3D projection. B. Arbuscules in cells. Arrowheads and dashed line depict the cell walls.

### 1.2 Critique of features analysed by Raman and micro-FTIR in Qu *et al.* (2015)

Qu *et al.* (2015) used Raman spectroscopy and micro-FTIR to analyse carbonaceous materials (CM) from Rhynie Chert fossils. Regarding the differences noted between fungi and plants and within plant tissue systems, the results should be treated with caution because some of the identifications appear to be erroneous.

1.2.1 The feature identified as a plant epidermis (Qu *et al.*, 2015; Fig. 3d, e, red dot; Fig. 9c) is acellular and far too thick (∼ 100 µm wide). It is a shrinkage feature that is commonly seen around the perimeter of plant stems at Rhynie (Trewin & Fayers, 2016). During fossilization, shrinkage of the plant axis can produce a void where the epidermis shrinks away from the initial silica coatings. The void created subsequently becomes filled with silica. Raman measurements of this feature are therefore recording CM in the background silica matrix.

1.2.2 The area of formless CM is misidentified as the phloem or phloem sap (Qu *et al.*, 2015; Fig. 3d, e, green dot; Fig. 6, i-l; Fig. 8c; Fig. 9c). In all the plants at Rhynie, phloem-like tissues are located towards the centre of the axis not at its periphery (Kerp, 2018). This feature is part of the outer cortex of the plant axis, and here it seems to be partly decomposed. Tissues in this region commonly contain endophytic mycorrhizal fungi, so the Raman measurements could be recording the degraded remains of a mixed assemblage of plant cortical cells and fungal hyphae.

1.2.3 The features identified as fungi, epilithic [sic] filamentous fungi, fungal tufts, or fungal rhizoids (Qu *et al.*, 2015; Fig. 3g, h, red and green dots; Fig. 6, e-h; Fig. 9b) are not fungi. They are plant rhizoid cells, which are common in the species found at Rhynie (Kerp, 2018).

